# Mathematical Model of Colorectal Cancer Initiation

**DOI:** 10.1101/2020.02.08.939603

**Authors:** Chay Paterson, Hans Clevers, Ivana Bozic

**Affiliations:** Department of Applied Mathematics, University of Washington, Seattle, WA; Hubrecht Institute, Utrecht, Netherlands

## Abstract

Quantifying evolutionary dynamics of cancer initiation and progression can provide insights into more effective strategies of early detection and treatment. Here we develop a mathematical model of colorectal cancer initiation through inactivation of two tumor suppressor genes and activation of one oncogene, accounting for the well-known path to colorectal cancer through loss of tumor suppressors *APC* and *TP53*, and gain of the *KRAS* oncogene. In the model, we allow mutations to occur in any order, leading to a complex network of incomplete mutational genotypes on the way to colorectal cancer. We parametrize the model using experimentally measured parameter values, many of them only recently available, and compare its predictions to epidemiological data on colorectal cancer incidence. We find that the reported incidence of colorectal cancer can be recovered using a mathematical model of colorectal cancer initiation together with experimentally measured mutation rates in colorectal tissues and proliferation rates of premalignant lesions. We demonstrate that the order of driver events in colorectal cancer is determined by the combined effect of the rates at which driver genes are mutated and the fitness effects they provide. Our results imply that there may not be significant immune suppression of untreated benign and malignant colorectal lesions.

## INTRODUCTION

Worldwide efforts to sequence cancer genomes over the past decade have provided insight into the number and type of genetic alterations implicated in tumor progression^1^. Recent work suggests that the acquisition of just three driver alterations^2^, often inactivation of two tumor suppressor genes and activation of one oncogene^1,3^, is sufficient to produce an invasive phenotype. In the case of colorectal cancer, the most significant driver events are inactivation of the tumor suppressor genes *APC* and *TP53*, and activation of the *KRAS* oncogene^4^. Activation of an oncogene requires a mutation of a single copy of the gene; inactivation of a tumor suppressor gene requires two genetic alterations (inactivation of both copies)^5^, leading to ~5 mutational stages on the way to colorectal cancer.

Early work on multistage modeling of colorectal cancer incidence aimed to determine the number of mutations (stages) needed for cancer initiation by fitting multistage models to cancer incidence data^6–8^. Notably, Armitage and Doll studied sequential accumulation of neutral mutations in a tissue, and reported ~6 stages were needed for colorectal cancer initiation^6^. Knudson used mathematical modeling and incidence data to conclude that two hits are needed for initiation of retinoblastoma; this finding subsequently being confirmed by the discovery of the Retinoblastoma (*Rb*) tumor suppressor gene^7^. Later works studied *n*-stage models where mutations accumulate in a pre-determined (linear) order, typically allowing the last pre-malignant stage to provide proliferative advantage to cells, with the goal to estimate mutation and proliferation rates from cancer incidence data^9,10^. However, even though some driver mutations tend to be acquired before others on the way to cancer, this order is frequently violated^11–13^.

In contrast to previous approaches, here we use a genomically-informed stochastic model in which we allow mutations to occur in any order, leading to a biologically-realistic complex network of possible evolutionary pathways that lead to cancer. Our approach is based on experimentally testable hypotheses about molecular mechanisms, and gives predicted incidence and timing of mutations as outputs, rather than looking backwards from epidemiological studies of incidence. We parametrize the model using recent experimental estimates of in vivo mutation rates in the colon^14^ and selective advantages of driver mutations in question^15,16^ and show that reported colorectal cancer incidence rate can be recapitulated by assuming that *APC* inactivation and *KRAS* activation provide small proliferative advantages to colorectal crypts, which is consistent with current experimental measures. We use the model to provide estimates of the relevant timescales and mutation order in the earliest stages of colorectal tumorigenesis.

### MODEL AND PARAMETERS

We develop and analyze a stochastic mathematical model of colorectal cancer initiation through acquisition of three driver events: inactivation of tumor suppressors *APC* and *TP53*, and activation of the *KRAS* oncogene. In this model, normal tissue consists of *N* colorectal crypts. Inactivation of a tumor suppressor gene (TSG) requires inactivation of both alleles. Each allele of a TSG can be inactivated through either mutation or loss of the allele. Loss of both alleles is not allowed, as this is usually fatal for the cell. An oncogene can be activated through mutation of a single copy of the gene. *r*_G_ is the rate of mutation of a gene G, and *r*_LOH_ is the rate of loss of an allele of a tumor suppressor gene (Methods). We do not impose any particular order in which these genetic events are acquired. This leads to a complex network of available 5-step pathways from an initially wild-type colorectal crypt to the one that has collected all three driver events (Fig. 1A).

**Figure 1.**
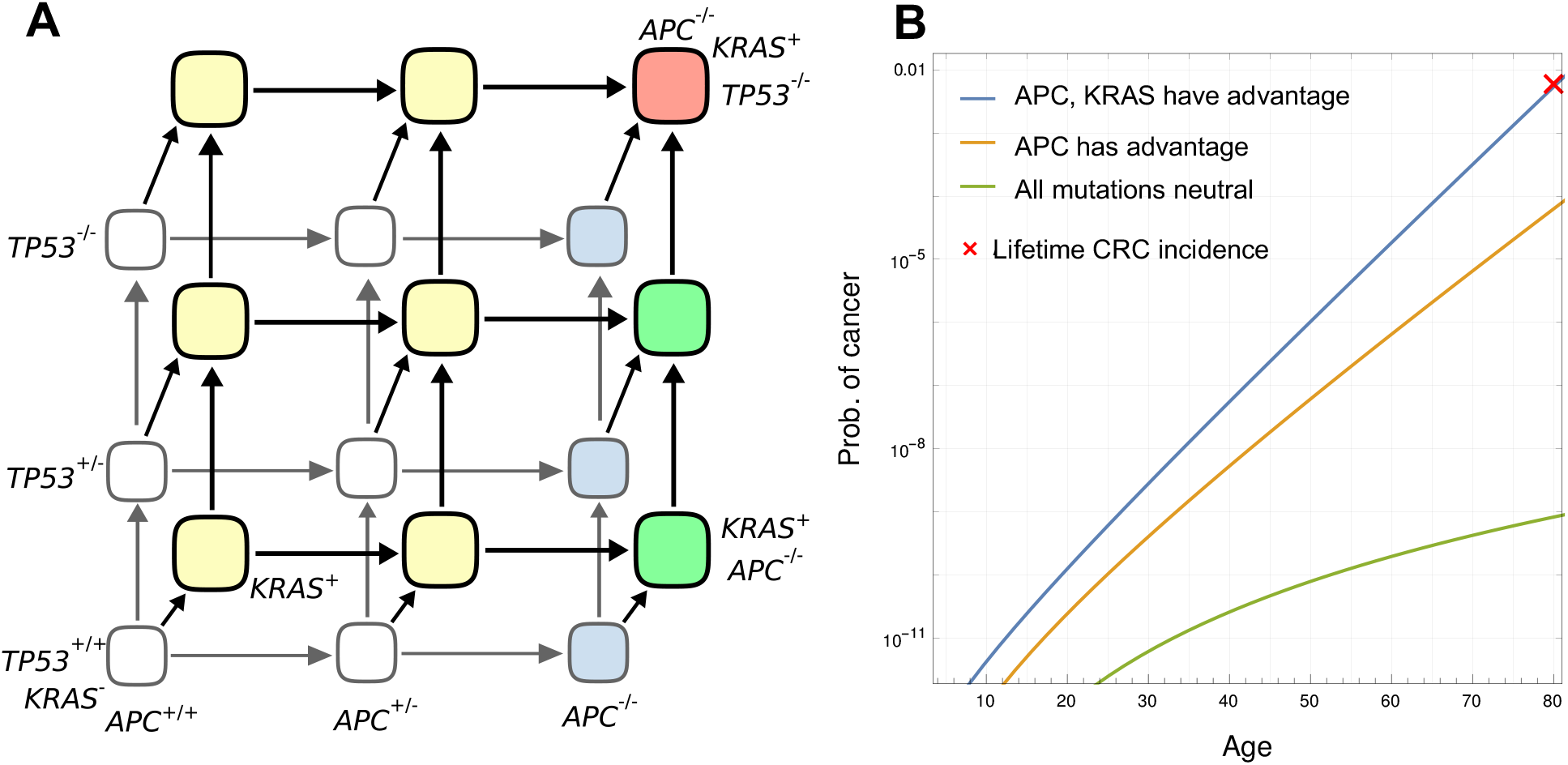
**A,** Schematic of colorectal cancer initiation. Healthy (wild type) crypt is in the lower left corner, and a fully malignant crypt in the top right. **B,** Comparison of Formulas (1), (2) and (3) and lifetime incidence of colorectal cancer containing mutations in *APC*, *KRAS* and *TP53*.

Many of the parameter values for this model have only recently become available. Mutation rates are gene-specific: they are the product of the number of positions in the gene that will result in (in)activation if mutated (driver positions), and the point mutation rate. The number of driver positions in the specific gene can be estimated from genetics knowledge and cancer mutation databases^17,18^ (see Methods for details). Intestinal stem cells reside at the bottom of intestinal crypts, with each crypt containing 5-8 stem cells^19^. It has recently become possible to measure the rate at which mutations accumulate in normal tissues, using organoids developed from single healthy stem cells from the colon, small intestine and liver^14^. Blokzijl and colleagues reported that individual stem cells of the colon accumulate about 40 mutations per year across the entire genome^14^, leading to a rate of *u* ~ 10^−8^ mutations per year per base pair. New mutations that appear in a single crypt will either be lost or will fixate in the crypt^20^, suggesting that the measured rate of accumulation of mutations applies at the level of a single crypt.

There are approximately *N*=10^7^−10^8^ crypts in the human colon^21,22^. Normal crypts divide only very rarely^16^, and we set their division rate to 0. It has been shown that *APC* inactivation and *KRAS* activation lead to clonal expansion of crypts carrying one of these alterations. Division rates of *APC*^−/−^ and *KRAS*^+^ crypts have been recently measured to be *b*_1_ = 0.2/year and *b*_2_ = 0.07/year, respectively^15,16^. In contrast, *TP53*^−/−^ is not thought to provide any fitness advantage on its own in normal conditions^23^.

### TIME OF APPEARANCE OF THE FIRST MALIGNANT CRYPT

We are interested in the time of appearance of the first crypt that has collected all three driver events – the first malignant crypt. Approximately 15% of colorectal cancers harbor all three driver events in question (*APC*, *TP53* and *KRAS*)^24^. The lifetime risk of colorectal adenocarcinoma^25^ is ~4.2%. Thus the lifetime risk of developing colorectal cancer containing these three driver events is ~0.6%. We will compare our model predictions to this incidence rate.

We analyze the model through computer simulations and analytic calculation. First, we study a model in which driver events are neutral until all three drivers are collected and show that this assumption leads to incidence rates that are many orders of magnitude lower than what is reported. In the case of neutral accumulation of driver mutations, in which all three drivers need to accumulate in a crypt in order for it to start a malignant expansion, the probability that a malignant crypt exists by time *t* is given by (Supplementary Methods)

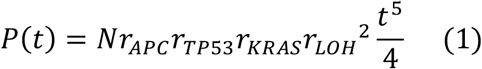

In Fig. 1B, we plot the probability of a malignant crypt as a function of time obtained using equation (1). If all mutations are neutral, probability of a malignant crypt at age 80 years is less than 1 in a billion – seven orders of magnitude lower compared to the ~1% of the population that develop colorectal cancer through acquisition of these three driver events. The assumption that all three driver mutations are neutral is thus not consistent with reported incidence, disagreeing by seven orders of magnitude.

Our results imply that at least one of the first two driver events, possibly both of them, have to provide the cells carrying them with a fitness advantage that allows them to increase in size by a factor of about 10^7^ (or alternatively, with an increase in mutation rate of comparable magnitude).

In agreement with this analysis, it has been shown that *APC* loss in colorectal stem cells does lead to a fitness advantage, and that stem cells that sustain *APC* loss will go on to produce an adenoma (polyp), which can be up to a centimeter in diameter or larger^26^. This will dramatically increase the chances for subsequent driver mutations within one of the cells of the *APC*-mutant clonal polyp. It was reported that *APC* inactivation leads to an increased rate of colorectal crypt fission, on the order of ~0.2/per year^15^.

We generalize equation (1) to the case when *APC* inactivation provides crypts with proliferative advantage in the form of division rate *b*_1_ (Supplementary Methods). In that case, probability of a fully malignant crypt being present by time *t* is given by

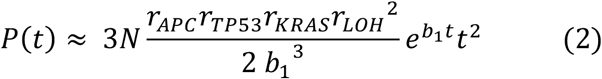

In Fig. 2A, we plot the probability of a malignant crypt as a function of time obtained from equation (2). Assuming that *APC* inactivation provides proliferative advantage to crypts leads to the probability of a malignant crypt at age 80: *P*(80) = 10^−5^, which is still ~3 orders of magnitude lower than reported incidence.

**Figure 2.**
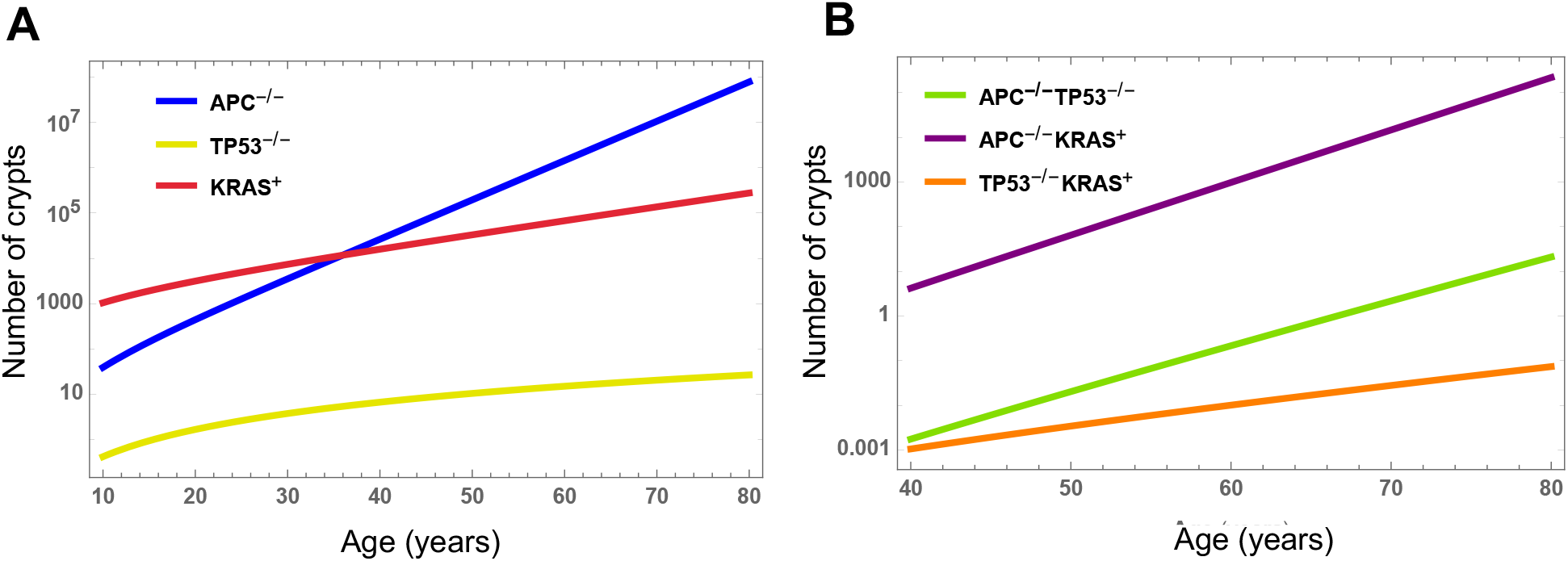
Expected number of single-(A) and double-mutant (B) colorectal crypts as a function of time. Formulas for expected numbers of mutated crypts used to produce the plots are given in Supplementary Methods.

We note that equation (2) does not take into account the increased chance that an *APC*-mutated stem cell will take over a crypt. It was measured that a stem cell with a single inactivated copy of the APC gene (APC^+/−^) has a 2.1 times higher chance to take over a crypt compared to a wild type stem cell, and that a stem cell with both APC copies inactivated (APC^−/−^) has a 2.8 times increased chance to take over a (APC^+/−^) crypt^23,27^. Thus the probability from equation (2) should be multiplied by *c*_1_=2.1*2.8=5.88 to obtain the corrected probability of a malignant crypt at age 80, *P_c_*(80) = 6*10^−5^, still two orders of magnitude lower than reported incidence.

Finally, we study a model in which *APC* and *KRAS*, but not *TP53* provide selective growth advantage to crypts, as *TP53* mutation does not seem to provide selective advantage on its own in normal conditions^23^ (i.e. without the presence of colitis). *KRAS* has been reported to provide growth advantage to colorectal stem cells and crypts, typically leading to small clonal expansions of *KRAS*-mutated crypts^28^. Recently the proliferation rate of *KRAS*-mutated crypts has been measured to be *b*_2_ = 0.07/year^16^. As before, we set division rate of *APC*-inactivated crypts to *b*_1_= 0.2/year, and assume crypts that have both *APC* inactivated and *KRAS* activated proliferate at rate *b*_12_ = *b*_1_ + *b*_2_ = 0.27/year. This amounts to assuming that there are no epistatic interactions between *APC* and *KRAS*: while this is an area of active research, the size and direction of any such effect, if it exists, is not presently known with confidence for these two genes^29^.

We derive the approximate formula for the probability that a malignant crypt is present by age *t* (Supplementary Methods):

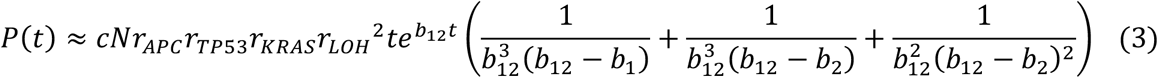

Here *c* = *c*_1_ * *c*_2_ = 5.88 * 3.6 = 21.2 is the correction to the formula to account for increased chance of fixation of APC and KRAS mutated stem cells in a crypt^23,27^. We plot equation (3) as a function of age *t* in Fig. 1B. In particular, the chance that a malignant crypt is present at age 80 is ~0.5%, very close to the lifetime risk of colorectal cancer through the *APC*/*TP53*/*KRAS* pathway of 0.6%. We also compared equations (2) and (3) to computer simulations of the mathematical model of colorectal cancer initiation. Equation (2), and equation (3) up to age ~60, are in excellent agreement with computer simulations of the process; equation (3) starts to overestimate the probability of a malignant crypt at later times (Fig. S1).

### NUMBER OF MUTATED CRYPTS

We next use our model to calculate the number of mutated colonic crypts as a function of time. The formulas for expected numbers of crypts that have inactivated a single tumor suppressor or activated the *KRAS* oncogene are given in Supplementary Methods and plotted in Fig. 2A. *KRAS* activation requires only a single hit; initially, the number of *KRAS*-mutated crypts is much higher than crypts with inactivated *APC* or *TP53*. However, due to the much faster fission rate of *APC*-inactivated crypts, their number surpasses the number of *KRAS*-mutated crypts by mid-age. The number of crypts with only *TP53* inactivated remains low throughout the human lifetime, due to a lack of growth advantage provided by *TP53* alone (Fig. 2A).

Formulas for the expected numbers of double-mutant colonic crypts as a function of age are given in Supplementary Methods and plotted in Fig. 2B. *APC*^−/−^*KRAS*^+^ crypts far outnumber the other two types of doubly-mutated crypts: by age 60, the expected number of *APC*^−/−^*KRAS*^+^ crypts is ~1000, whereas the numbers of *APC*^−/−^*TP53*^−/−^ and *TP53*^−/−^ *KRAS*^+^ crypts stay below ~10 and below 1, respectively, throughout the human lifespan (Fig. 2B).

### ORDER OF MUTATIONS

The primary outstanding question that we aim to address is: in what order do these three functional mutations typically appear over the course of cancer initiation? To answer this, we must differentiate between two relevant questions: (i) in which order do driver mutations arise in a typical colorectal crypt, and (ii) what is the order of mutations that was experienced by the first malignant crypt? A typical colorectal crypt will not have any of the driver events in question: 99.998% of crypts will not have activated *KRAS* or deactivated *APC* or *TP53* by age 80 (Supplementary Methods). Note that here we take into account only the fraction of *N* initial healthy crypts that have accumulated a driver event, and not the additional crypts that derive from abnormal division of crypts that have accumulated drivers. Of the small number of crypts that have collected a driver event, the most likely first event in a typical crypt is the activation of *KRAS*, as it only requires a single mutation (hit). This starts to answer question (i): large majority of crypts will be functionally normal, and the most likely first driver accumulated by a typical colorectal crypt is *KRAS*.

We are however, more interested in question (ii), the order of driver events that will actually produce a malignant crypt during a human lifespan. To answer this question, we developed a new Monte Carlo simulation, related to the previous simulation we used to measure cancer incidence. In the new simulation, instead of simply moving cells between different sites in the “space” of possible genotypes, all possible orders of the five mutational steps were tracked (Supplementary Methods). Cells could arrive at one of the four possible cancerous genotypes (corresponding to double mutation, or mutation and LOH of the two tumor suppressors) by 270 different possible paths (Fig. 3A left). The probability for each of these paths to be taken by a crypt that became malignant by age 80 years was then measured, and that path was classified into one of the six possible sequences of the three driver events (Fig. 3A right).

**Figure 3.**
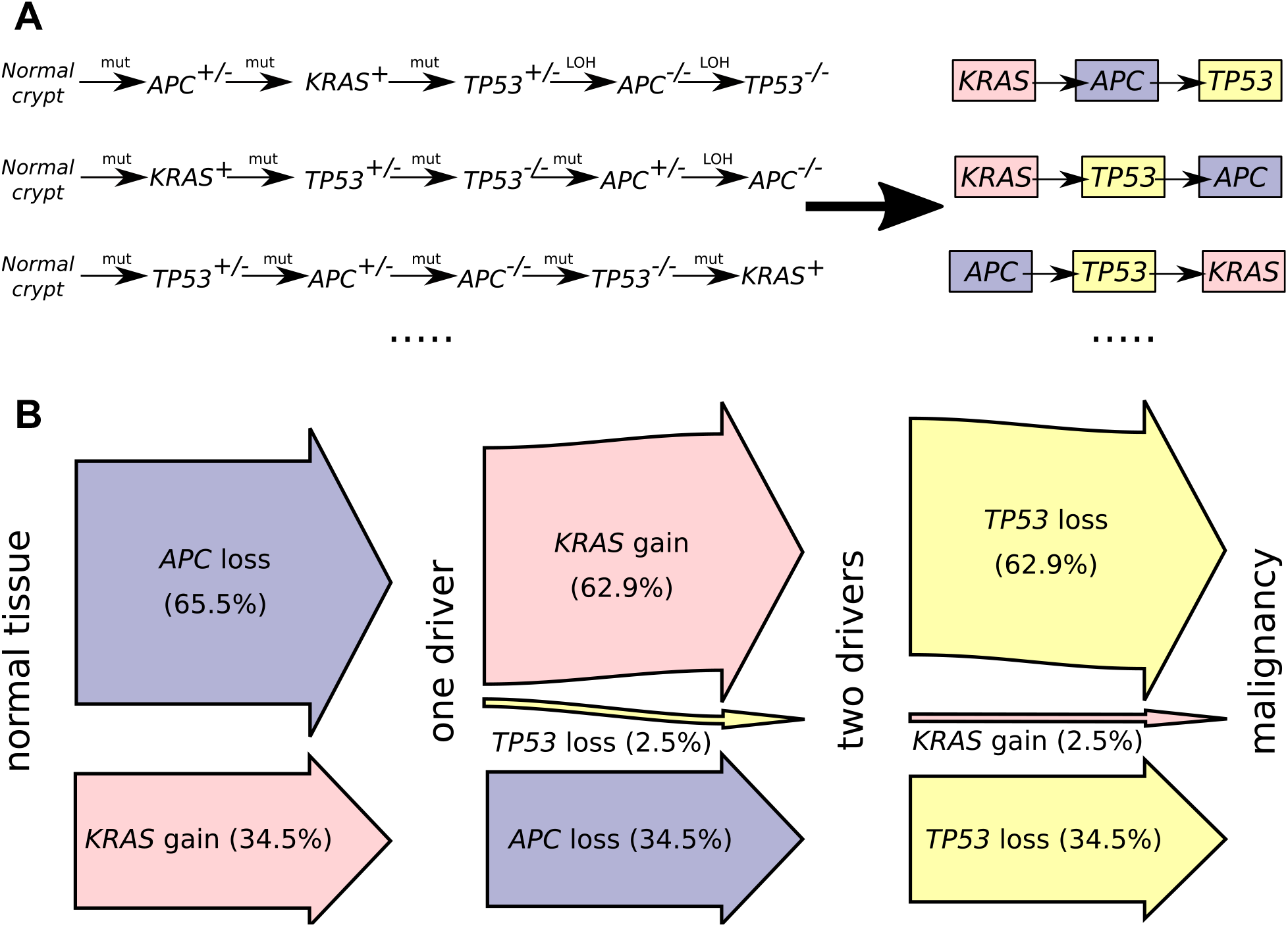
Order of acquisition of driver mutations in the mathematical model of colorectal cancer initiation. **A,** Three example 5-step paths (out of 270 possible paths) that a crypt can take in order to become malignant. We classify each path that leads to malignancy within 80 years in our computer simulation into one of the six possible orderings of the three driver mutations. **B,** Likelihood of driver mutation order on the path to colorectal cancer, obtained from computer simulation of our mathematical model.

The result of this mutational order analysis is shown in Figure 3B. The majority of the first driver events in a path that ended in malignancy are in fact the inactivation of *APC*, accounting for 65.5% of first events. All of the remainder (34.5%) were gain of *KRAS*, as *TP53* was never observed to be lost first. The majority of second driver events were gain of *KRAS* (62.9%), all of which followed from *APC* being lost first. The second most common second driver event (34.5%) was the inactivation of *APC*: in a small minority of cases (2.5%), *TP53* was lost second. The last driver event in an outstanding majority of cases (97.4%) was the loss of *TP53*.

It is remarkable that the loss of *APC* was so often the first mutational event to have occurred on the path to malignancy, given that in a typical colorectal crypt, *KRAS* gain is the first driver event. Our explanation for this is that the loss of *APC* carries the largest fitness advantage of any event in our model, so cells arriving at a cancerous genotype will tend to come from lineages in which *APC* is lost early. The loss of *TP53* was overwhelmingly the last event to occur out of the three: we attribute this to the lack of a fitness advantage ascribed to *TP53* loss, and the consequent small number of crypts which carry *TP53*^−/−^ as one of the first driver events (Fig. 2A,B).

The order of mutations in our mathematical model is surprisingly similar to the well-known order of mutations on the path to colorectal cancer that was first described by Fearon and Vogelstein^30^. In line with our results, *KRAS* mutations were found to be present with high frequency in the earliest preneoplastic lesions in the colon, and absent in morphologically normal crypt areas^31^. However, while *KRAS* mutations are very common in small nondysplastic lesions, such lesions have limited potential to progress to larger tumors and ultimately cancer^32^.

In contrast, lesions with *APC* inactivation as the first event have a much stronger capacity to progress^32^. *APC* was found in the earliest neoplastic lesions that can be examined^32^, and is believed to precede both *KRAS* and *TP53* in the adenoma-carcinoma sequence of colorectal cancer^30^. Progression of *APC*-mutated adenomas usually occurs through subsequent acquisition of *KRAS* and other genes^32,33^. *KRAS* mutations were found in ~10% of colorectal adenomas smaller than 1cm, and in ~40% of adenomas larger than 1cm^34^, thus most likely to appear after *APC* was inactivated and started an adenoma, but before malignant transformation. As in our model, evidence points to inactivation of *TP53* being a relatively late event in colorectal tumor progression^11,30,35^, associated with the transition from benign adenoma to malignant carcinoma^36^.

## DISCUSSION

In this paper we develop and analyze a stochastic model of colorectal cancer initiation through a biologically-realistic complex network of possible mutational genotypes on the path to malignancy. In this model, we calculate the probability that a colorectal crypt has collected three driver events of interest (*APC*, *KRAS* and *TP53*) and became malignant as a function of age. We find that the reported incidence of colorectal cancer can be recovered using a mathematical model of colorectal cancer initiation together with experimentally measured mutation rates in colorectal tissues and proliferation rates of premalignant lesions. This result has several important implications. First, it sheds light on the important mutation- selection debate in cancer, showing that neither increased mutation rates nor large proliferation rates of premalignant lesions are necessary to explain observed cancer incidence rates. Another intriguing implication of our findings is that the immune system may not be exerting significant pressure on the pre-malignant and malignant lesions during tumor evolution, as the reported incidence of colorectal malignancy (which includes those cancers that escaped immune surveillance) is not lower than the incidence of malignant crypts predicted by the model. This finding is in agreement with recent reports that there may not be significant immune selection pressure during natural tumor evolution^37^.

Our model provides a precise quantitative understanding of the process of premalignant and malignant transformation in the human colon, and can be used as a starting point in evaluating early detection and treatment efforts. In particular, our results imply that, even though individual *KRAS*^+^ lesions are less likely to progress to malignancy compared to *APC*^−/−^ lesions, *KRAS* mutation may be the initiating driver event in as many as a third of colorectal cancers containing this alteration; thus efforts to target *KRAS*^38^ may prove particularly important in a significant fraction of patients.

Many previous models of colorectal cancer initiation study dynamics of individual cells in colorectal crypts^39^. We circumvent the need to analyze the accumulation of mutations in the stem cell populations of colonic crypts by using the recently measured rate of accumulation of mutations in normal tissues^14^. New mutations are either lost or fixate in the crypt^20^, which allows us to focus on crypts as the level of selection. Research suggest clonal stabilization time in mice crypts is ~1 month and in human colon ~1 year^20,40^. As the reported times apply to neutral mutations, it is expected that driver mutations that we are studying fixate even faster, on the order of months if not less, and thus the measured rate of accumulation of mutations in colonic stem cells can also be applied at the level of individual crypts.

Healthy cells are chromosomally stable, but most colorectal cancers are chromosomally unstable. Thus at some point during tumor evolution cells acquire chromosomal instability (CIN). Several early studies suggested that CIN is an early event in colorectal tumorigenesis, even preceding *APC* inactivation^41,42^. However, more recent studies suggest that extensive aneuploidy such as seen in the presence of CIN occurs after inactivation of *TP53*^43,44^ or possibly inactivation of both *APC* and *TP53*^45^. Since *TP53* inactivation usually follows *APC* inactivation and *KRAS* activation^30,35^, it is likely that acquisition of our three driver events of interest precedes significant CIN.

We use the measured accumulation rate of mutations in normal tissues, which includes contribution from the mutations that arise from unavoidable errors associated with DNA replication, as well as any environmental factors^46^. Once the relative contributions to the overall mutation rate from these two sources are known, our model can be used to quantify the relative proportions of sporadic colorectal cancers that arise predominantly due to replicative errors (“bad luck”)^21^ or environmental factors^47^.

We find that only a small fraction of human colorectal crypts will carry one of the driver mutations we study (fully inactivated *APC* or *TP53*, or activated *KRAS*). Our results do not contradict the recent report of probable driver mutations in around 1% of normal colorectal crypts in middle-aged individuals^48^. Lee-Six et al.^48^ examined ~1000 colorectal crypts from 42 individuals and examined the crypts for mutations in 90 known colorectal cancer genes. None of the crypts carried a driver mutation in *KRAS*, and no crypts carried two inactivating alterations of *APC* or *TP53*, implying that the frequency of such alterations is <0.1%.

Here we assume that point mutation rate does not increase during early stages of pre-malignant colorectal neoplasia. As more detailed experimental measures become available, our framework could in the future be extended to account for the potential increase in mutation rates of incomplete mutational genotypes on the road to malignancy. Of note, as the predicted CRC incidence from our model is of the same order of magnitude as the reported incidence, we do not expect mutation rates to be significantly increased prior to malignant transformation^49^.

## METHODS

### Inactivation rates for TSGs

A single copy of a wild-type TSG can be inactivated through either mutation, or loss of the allele via a loss of heterozygosity event (LOH). Inactivating mutation rate per two wild-type copies of a tumor suppressor gene is *r*_TSG_, where TSG can be either *APC* or *TP53*. Rate of LOH per two wild-type copies of a TSG is *r*_LOH_. Thus, if the crypt is TSG wildtype (TSG^+/+^), it can inactivate the first copy of the tumor suppressor through either mutation or LOH with a joint rate of *r*_TSG_ + *r*_LOH_. If one copy of a TSG is already inactivated (TSG^+/−^) through mutation, the second copy can be lost by either mutation or LOH, with the rate (*r*_TSG_ + *r*_LOH_)/2. If, on the other hand, the first copy was lost through an LOH event, the second copy can only be lost through mutation, with rate *r*_TSG_/2. In other words, we are implicitly assuming that the loss of both copies through LOH is fatal for the cells.

### Gene-specific driver mutation rates

We estimate mutation rates of individual driver genes *D*, *r*_D_, by multiplying the base pair mutation rate *u* with the number of positions in the gene that will lead to (in)activation if mutated, *n*_D_. In other words, *r*_D_ = *n*_D_\**u*. We previously analyzed all possible mutations at each of the 8,529 protein-coding bases in the *APC* gene^17^. Our analysis found that there are *n*_APC_=604 driver positions in the *APC* gene, which will result in allele inactivation if mutated. There are 1179 protein-coding bases in the *TP53* gene. Assuming that the *TP53* statistics follows that of the *APC*, there are *n*_TP53_=1179*604/8529=83 driver positions in the *TP53* gene. Finally, there are approximately *n*_KRAS_~20 position in the *KRAS* gene that lead to its activation if mutated^50^.

### Rate of LOH

The rate of LOH in the absence of chromosomal instability can be estimated from mutational data from colorectal cancers with microsatellite instability (MIN), since MIN and CIN are mutually exclusive. Huang et al.^51^ report that the ratio of MIN cancers with two inactivating mutations in *APC* to those with one inactivating mutation and one LOH event is 1:7. Let us denote by *k* the ratio between the rate of LOH and the rate of inactivating mutation in MIN cancers. The fraction of cases where first event is LOH is *k*/(*k*+1), and the fraction of cases where first event is mutation is 1/(*k*+1). The fraction of cases where first event is a mutation and second event is LOH is 1/(*k*+1) *k*/(*k*+1), and the fraction of cases where both events are mutations is 1/(*k*+1) *1*/(*k*+1). It follows that the ratio of cases where one event is LOH to those where both events are mutations is *k*^2^ + 2*k*. From *k*^2^ + 2*k* =7 we obtain that *k* ~ 1.8. Thus the rate of LOH at the *APC* locus in MIN cancers is 1.8 times higher compared to the rate of inactivating mutation. Colorectal cancers with MIN have ~10 times increased point mutation rate compared to non-MIN cancers^52,53^, so we conclude that the rate of LOH is ~18 times higher compared to the normal rate of inactivating mutation in *APC*: *r*_LOH_ = 18* 604*10^−8^ ~ 10^−4^ per year. We assume that rates of LOH at *APC* and *TP53* are approximately the same.

## Supplementary Methods

### Basic model

We study a stochastic model of accumulation of oncogenic mutations in individual crypts of the colon. We specifically focus on mutations in three genes: *APC*, *KRAS* and *TP53*. Initially, there are N wild-type crypts. A wild-type crypt genotype is denoted by (0,0,0), corresponding to no mutations in any of the three genes. A wild-type crypt can lose a single copy of the *APC* gene via LOH leading to genotype (1,0,0). If the first copy of *APC* is lost via mutation, this leads to genotype (2,0,0). If both copies of APC are lost, the corresponding genotypes are (3,0,0) if one copy is lost via mutation and one via LOH, or (4,0,0) if both copies are lost through mutation. Loss of both copies via LOH is not allowed. Rates at which these events (mutations and LOH) occur in individual crypts are described in Methods. Genetic alterations in *TP53* occur in a similar manner as described above for *APC* (although with different rates). For example, we denote by (0,2,0) a crypt that has mutated a single copy of *TP53*. Genotype that denotes that *KRAS* is mutated is (0,0,1). The assumptions described above lead to 50 possible genotypes (5 5 2), including the four malignant genotypes (3,3,1), (4,3,1), (3,4,1) and (4,4,1).

Crypts that have accumulated certain driver alterations, such as mutation in *KRAS* or loss of both copies of *APC*, undergo fission, and increase in number. We assume that this increase occurs stochastically with some growth rate determined experimentally (see main text). We can therefore view the model as studying the movement of crypts through a “space” consisting of different possible genotypes, and “birth” (fission) of crypts on certain “sites” with an advantage. When a mutation occurs, a crypt moves from one of these sites to another; when fission occurs, the number of crypts on an advantageous site increases by 1.

### Analytical approximation

For the purposes of our analytical approximation, we assume that double mutations do not occur. This should be true as long as the rate of loss of heterozygosity is much less than the point mutation rate for each gene: since *r*_*LOH*_ ≈ 10^−4^/*yr* and the base point mutation rate *u* = 10^−8^/site/yr, this assumption will be accurate for genes with significantly fewer than 10^4^ sites at which driver mutations can occur. This reduces the number of distinct genotypes we need to consider to 32, with the only malignant genotype being (3,3,1).

We use the mean-field approximation^54^ for the probability *P(t)* that at least one cancerous crypt is present by time *t*:

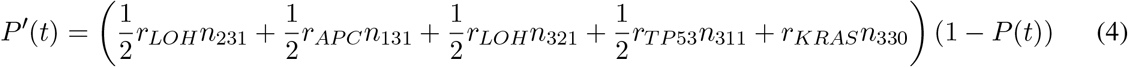

Here *n*_*ijk*_ = *n*_*ijk*_*(t)* denotes the expected number of crypts of genotype *(i, j, k)* at time *t*. Solution to equation (4) can be written

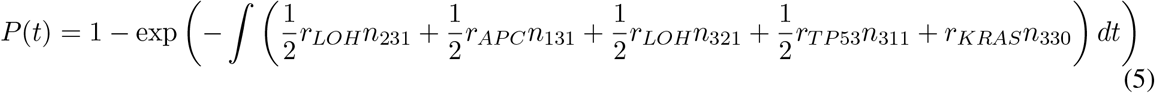

In the limit that *rt* ≪ 1, for all mutation rates r, we can approximate expected numbers of crypts with non-malignant genotypes using the solutions to the following system of equations:

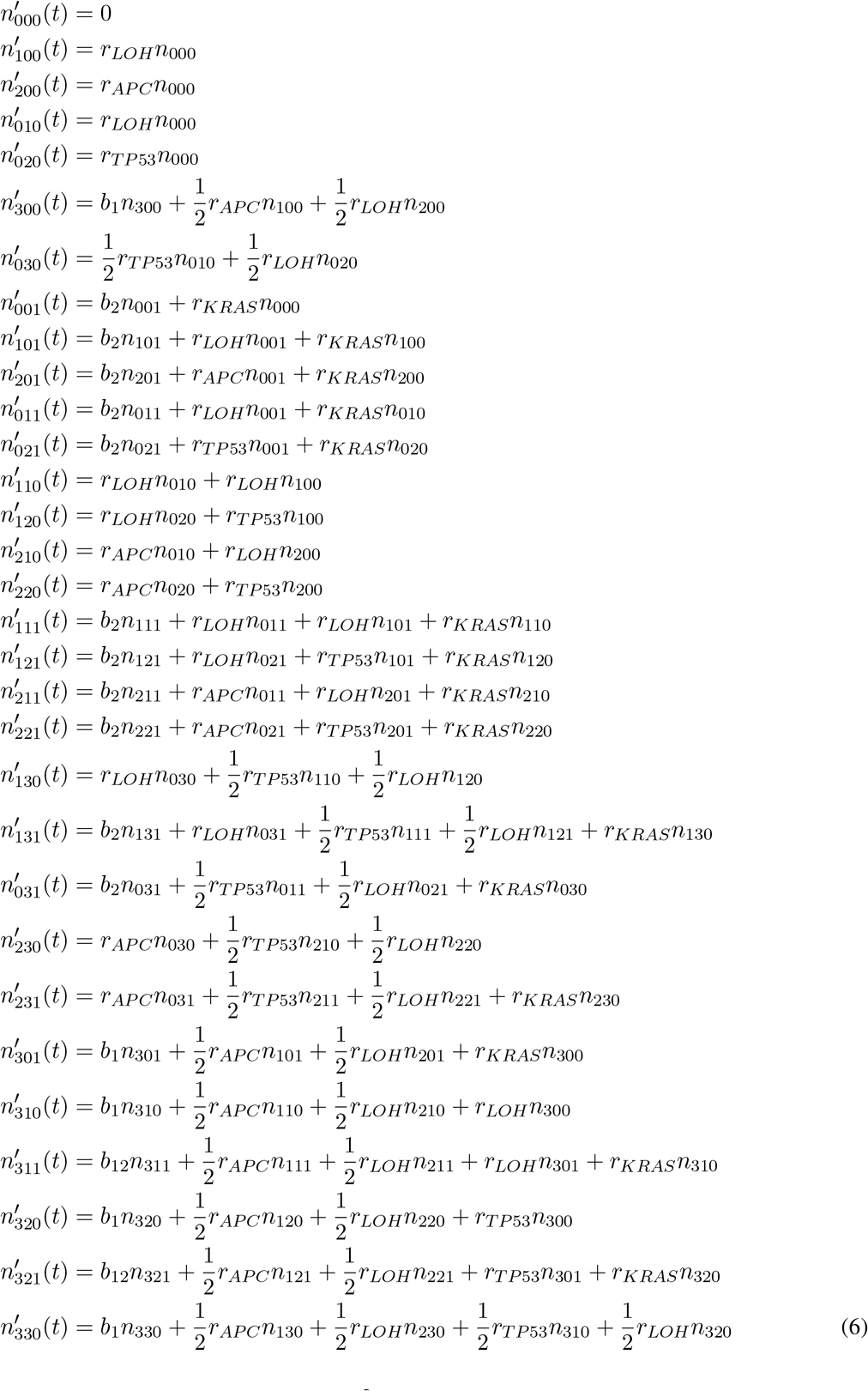

In the above system, we have neglectied the negative (“outflow”) terms. Since the largest mutation rate, *r*_*LOH*_ = 10^−4^/*yr*, and the largest time *t* = 80*yr*, it will always be the case that

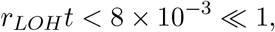

and this approximation is fair. The relevant initial conditions are that all crypts are initially wild-type (i.e. normal tissue), thus:

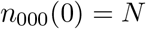

and there are no crypts of any mutant genotype, so for all other ijk:

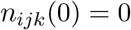

As the system (6) is linear, it can be efficiently solved using integrating factors.

### All drivers neutral

When *b*_1_ = *b*_2_ = *b*_12_ = 0, and *r*_*LOH*_*t* ≪ 1, we can note that

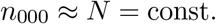

and all other solutions *n*_*ijk*_ to the approximate system (6) are simply polynomials in *t* (this follows from repeated integration of *n*_000_ = const.). In particular,

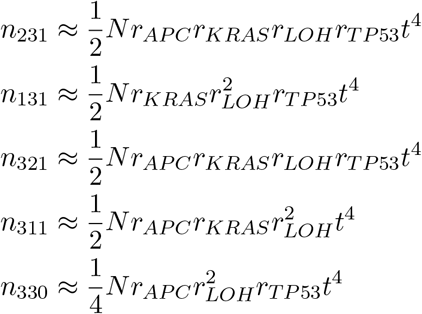

which together with (4) imply that

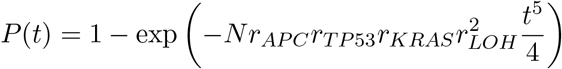

When *P* ≪ 1, this can be approximated as

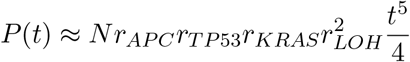

which is just equation (1) from the main text, analogous to Armitage & Doll’s famous result^6^.

### APC only has advantage

When *b*_1_ > 0 but *b*_2_ =0 and *b*_12_ = *b*_1_, we again have

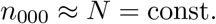

but terms corresponding to *APC*^*−/−*^ are different. This affects the calculation of *n*_321_, *n*_311_, and *n*_330_:

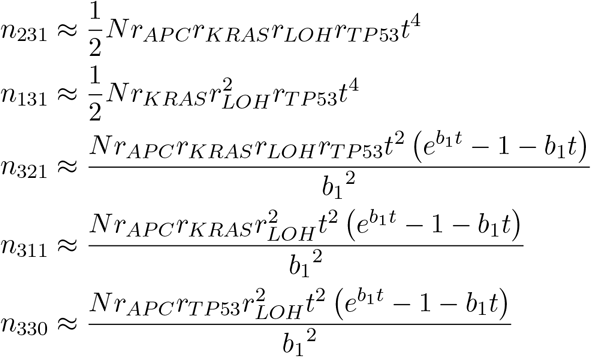

which implies that

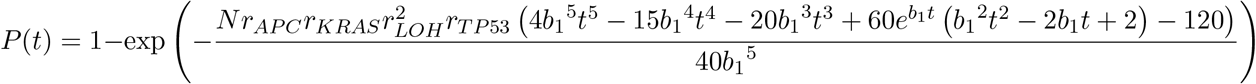

and, in the limit that P ≪ 1,

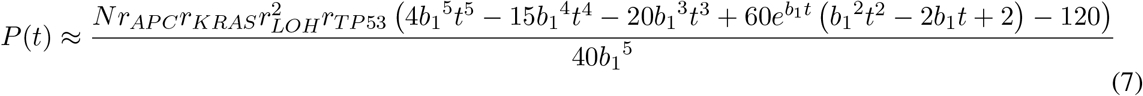

This can be simplified further with relatively little loss of accuracy (in the limit that *r*_*LOH*_ /*b*_1_ is small and *b*_1_t is large). The fastest growing term in (7) is 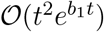: so, when *b*_1_*t* ≫ 1, this term must dominate all the others, and it should be the case that

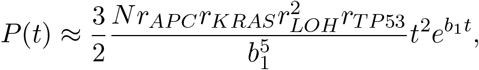

which is just equation (2) from the main text.

### APC and KRAS both have advantage

When *b*_1_ > 0, *b*_2_ > 0 and *b*_12_ > 0, the expressions for *n*_231_, *n*_131_, *n*_321_, *n*_311_, and *n*_330_ can be obtained by solving the system (6), from which *P(t)* can be found using expression (5). The expression for *P (t)* would take more than a page to write out, so we do not specify it here. However, despite its formidable appearance, given some reasonable hypotheses about *b*_1_, *b*_2_ and *b*_12_, the dominant term can be extracted relatively easily. We know experimentally that

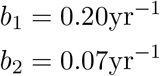

and have further postulated that *b*_12_ = *b*_1_ + *b*_2_. Under the weaker hypothesis that *b*_12_ > *b*_1_ > *b*_2_, then when (*b*_12_ − *b*_1_)*t* ≫ 1 (which given the above boils down to 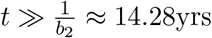), the dominant term will be

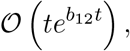

leading to

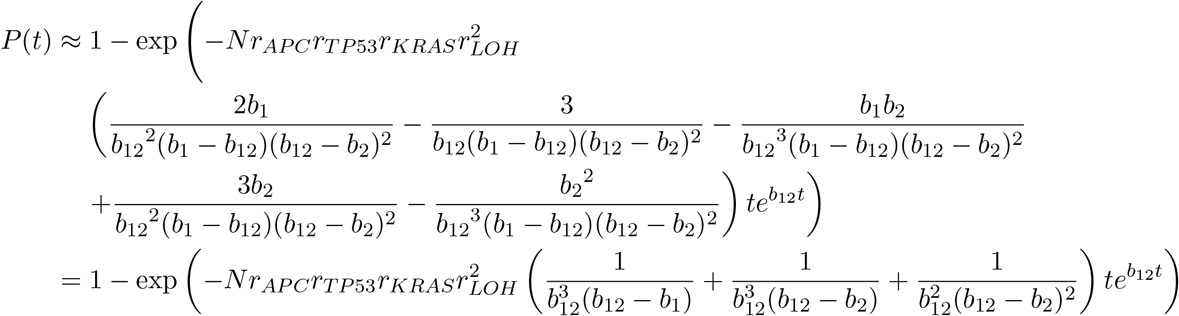

When *P* ≪ 1,

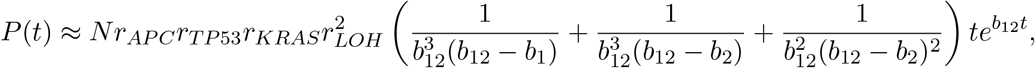

which up to a factor of *c* is equation (3) from the main text.

### Number of mutated crypts

Expected numbers of mutated crypts can be obtained using system (6). The expected numbers of crypts that have inactivated a single tumor suppressor or activated the KRAS oncogene are given by:

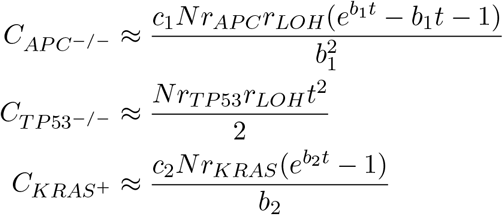

Expected numbers of double-mutant colonic crypts as a function of age are given by the following equations:

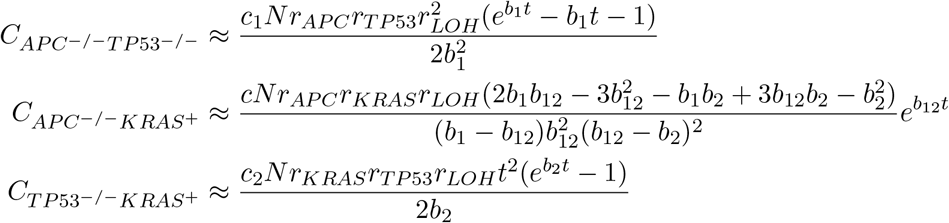

### Numerical simulations

For our numerical simulations, in contrast to the earlier analytical approximations, we make no simplifying assumptions regarding possible genotypes and mutations. Genotypes in which two successive mutations have occurred on some tumor suppressor gene are included in the space of all possible genotypes, even though they are much less frequent than genotypes where one copy is mutated and the other is lost.

Our first sets of simulations were performed with Gillespie’s Stochastic Simulation Algorithm^55^. This is an important test because the SSA provides an exact sample of the underlying master equation governing the random processes of our model. Let 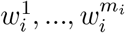 be the rates of possible events (mutations, LOH, fission) at site i. The weighted rate of an event is the product of the number of crypts at a site and the rate of the event in question: 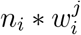.

In summary, the algorithm consists of the following:

1. Calculate the aggregate event rate Γ. This is essentially just the weighted sum of populations *n*_*i*_ at sites *i* and the overall event rate for that site, 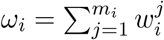:

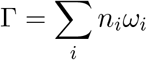
2. Sample an event according to a categorical distribution where the probability of each event is proportional to its weighted rate.
3. Draw time to next event Δ*t* from an exponential distribution with mean Γ^−1^.
4. Update the population according to the sampled event, advance time *t* by Δ*t*, and repeat steps 1-4.

To estimate probabilities of a specific event (such as cancer initiation, for example) having occurred by a given time *t*, many independent replicates *R* of the same process are obtained by the above method, the number of replicates *Q* in which the event occurred is noted, and the probability *P* is estimated as

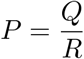

Clearly, probabilities significantly smaller than 1/R cannot be measured, and there will be some degree of stochastic noise in *P* scaling approximately with 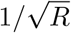 for small *P*. The accuracy of this (or indeed any similar) method of using stochastic simulations to estimate *P* is therefore limited by the number of replicates R that are practically achievable with the algorithm on the available hardware.

### τ-leaping

Although it is “exact” in the sense that it provides an exact sample of the underlying master equation, the Gillespie algorithm is computationally very expensive. To efficiently study rare events in the same framework as relatively common stochastic processes without making any simplifying assumptions that treat one process as deterministic – for example, to study mutations in the same framework as crypt fission, a much more common event – many replicates are required.

A more efficient *approximate* method of simulating stochastic processes was also developed by Gillespie, and is usually known as τ-leaping^56^. τ-leaping is closely related to Euler integration, in that the time step τ is held fixed, and time *t* advanced by this constant amount in each step of the simulation. The outline of the algorithm is as follows:

1. For each site i with population *n*_*i*_ and mutation event *i* → *j* with rate *r*_*i→j*_, generate a Poisson-distributed random variable Δ*n*_*i→j*_ with mean *n*_*i*_*r*_*i→j*_ τ. i.e.

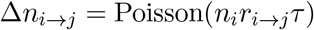 If *n*_*i*_*r*_*i→j*_τ is of a similar order of magnitude to *n*_*i*_, one should instead take Δ*n*_*i→j*_ to be max(Poisson(*n*_*i*_*r*_*i→j*_τ, *n*_*i*_) to ensure that the population of no site is ever negative. This is one among several possible operational choices^57^.
2. Move Δ*n*_*i→j*_ crypts from site *i* to site *j*.
3. For each site i with population *n*_*i*_ and fission rate *b*_*i*_, generate a negative-binomially-distributed random variable Δ*n*_*i*_ with mean *n*_*i*_*b*_*i*_τ.
4. Add Δ*n*_*i*_ crypts to site *i*.
5. Advance time *t* by τ, and repeat steps 1 to 4.

The choice of a negative binomial distribution instead of a Poisson distribution in step 3 reflects the “contagious” nature of the fission process, entailing exponential growth, and provides a faster convergence to the SSA as τ decreases^57^

We first verified that the results of the τ-leaping algorithm converged to the SSA as τ decreased, before replicates were gathered at a compromise value of τ = 0.01yr, which provided acceptable agreement (within observed stochastic errors in the SSA). This allowed 25-100 times as many replicates to be collected as in the pure SSA simulations, significantly increasing the accuracy with which probabilities P could be estimated.

### Lineage tracking simulations

Simulations that only track the number of crypts present on any given “site” do not carry information about the sequence of steps that any individual crypt took to get there. In other words, they cannot be used to enumerate the lineages of mutated crypts, or the order of mutations that occurred in the initiation of a typical cancer. We modified our simulations in order to address these questions as well.

The basic picture of having individual crypts move around a space consisting of “sites” and connections between these sites can remain in place, along with the algorithms used to study stochastic processes occurring on this space, as long as the interpretation of these sites changes somewhat. Instead of a “site” corresponding to one genotype, a “site” should now correspond to an ordered pair consisting of a genotype with a sequence of genotypes leading up to it. In other words, a “site” *ℓ* in this new structure is the ordered pair

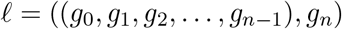

with g_*n*_ the most recent genotype visited in the sequence, where the crypts that have taken this pathway actually reside. This way, the different paths taken to the same g_*n*_ from g_0_ (the wild-type genotype) can be distinguished by the different g_1_, g_2_,…, g_*n* − 1_ in the sequence before g_*n*_. For clarity, we will refer to such “sites” as used in this set of algorithms as “lineages”. So, a “lineage” *ℓ* consists of a terminal genotype g_*n*_ where some population of crypts resides, along with a historical sequence (*g_0_,…, g_*n*−1_*) leading up to g_*n*_.

Two lineages, say *ℓ* and *ℓ*′, are connected if *ℓ*′ can be reached from *ℓ* by mutation or LOH. Writing out *ℓ* and *ℓ*′ termwise and comparing:

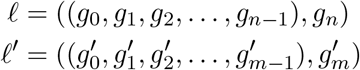

it should be clear that *ℓ* can be reached from *ℓ* if 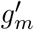 contains one more alteration than g_*n*_, and the paths leading up to g_*n*_ and 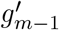 are identical. That is,

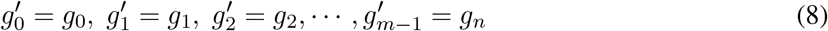

and the mutation rate

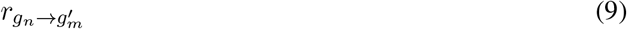

is nonzero.

Taken altogether, this describes a new graph structure, the space of possible lineages, derived from the previous space of possible genotypes. Once this space is constructed iteratively using rules (8) and (9), fundamentally similar algorithms can be used to simulate stochastic processes occurring on this space as on the space of possible genotypes. For practical reasons, we used tau-leaping with a τ = 0.01yr.

Due to the combinatorial nature of the paths taken, there are many more possible lineages than there are genotypes. There are 270 lineages that terminate in a cancerous genotype, for example: 120 involve two LOH events, a further 120 involve only one LOH event, and a final 30 involve no LOH events, and only double mutations. As expected, these last 30 are exceptionally rare, and the majority involve two LOH events, one for each tumor suppressor gene involved (*APC* and *TP53*).

**Figure S1.**
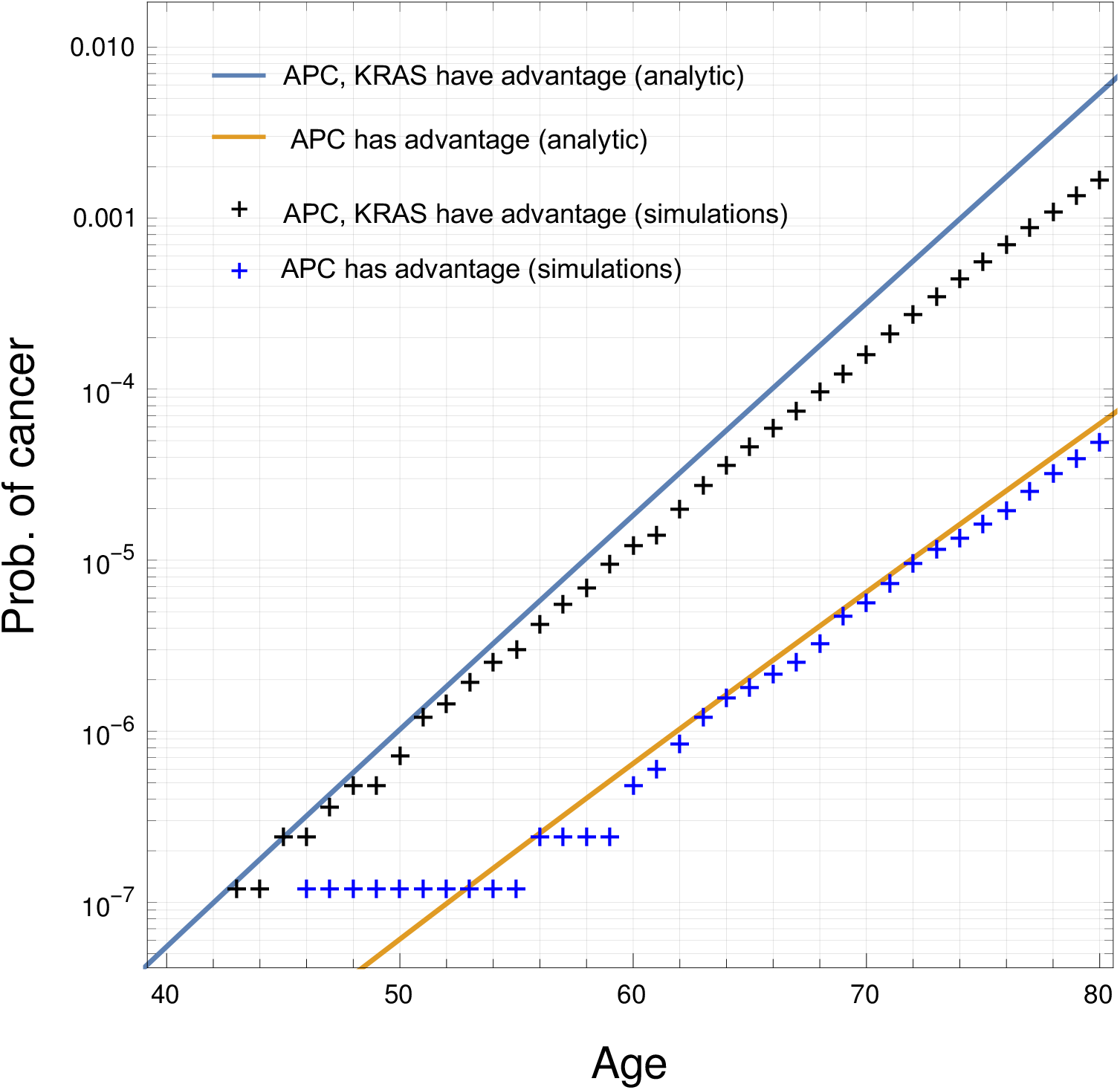
Comparison of analytic approximations and computer simulations for probability of a malignant crypt. Comparison of formulas (3) and (2) from the main text to τ-leaping simulations of the process of colorectal tumor evolution. 8. 10^6^ runs were performed for each of the two parameter combinations.

